# Study on endogenous inhibitors against PD-L1: cAMP as a potential candidate

**DOI:** 10.1101/2022.07.12.499690

**Authors:** Qiuyang Huang, Xiaoling Zang, Zhiwei Zhang, Xin Zhang, Mustafa R. K. Ali, Zhihua Lv

## Abstract

The discovery of new anticancer drugs targeting the PD-1/PD-L1 pathway has been research hotspots. In this study, a combination of biological affinity ultrafiltration (BAU), UPLC-HRMS, molecular dynamic (MD) simulations and molecular docking methods were applied to search for endogenous active compounds that can inhibit the binding of PD-L1 and PD-1. We screened dozens of potential cancer related endogenous compounds. The results showed that cyclic adenosine monophosphate (cAMP) had a direct inhibition effect on the PD-1/PD-L1 binding with an in vitro IC^50^ value of about 2.7 µM determined by homogeneous time-resolved fluorescence (HTRF) assay. The binding mode analyses for the cAMP - dimeric/monomeric PD-L1 complex indicated that cAMP was likely to bind to the dimeric PD-L1, since the binding free energies of the cAMP - dimeric and monomeric PD-L1 complex were about 23.6 and 15.1 kcal/mol, respectively, from MD simulations. The direct binding assay using surface plasmon resonance (SPR) method showed that cAMP could also bind to monomeric PD-L1 fixed on the sensor chip surface with a K_D_ value of about 1.72 mM. Our findings suggested that cAMP may directly inhibit the PD-1/PD-L1 interaction.

## Introduction

Cancer cells can escape immune surveillance through activation of inhibitory immune checkpoint proteins including PD-1, CTLA-4, LAG3, TIM3, TIGIT and BTLA^1^. Immune checkpoint blockade therapies have shown significant success on multiple malignancies and have provided substantial clinical benefits. Programmed death-1 (PD-1) expressed on activated T-lymphocytes has two ligands, programmed death-ligand 1 (PD-L1) and PD-L2, which are expressed on dendritic cells, macrophages, activated B and T cells, and mesenchymal stem cells, etc^2^. Also, PD-L1 is known to be markedly enriched on the cancer cell surfaces of many tumor types^3^. The binding between PD-L1 and PD-1 can inhibit T-cell proliferation and induce T-cell apoptosis, anergy and functional exhaustion^4^. Targeting the PD-1/PD-L1 pathway with monoclonal antibodies (mAbs) is an effective approach for cancer immunotherapy and it has been extensively studied and developed for treatment of multiple malignancies^4, 5^. To date, avelumab and other mAbs targeting PD-L1 have been approved by the US FDA^4, 6^. Very recently, Cercek et al. performed phase 2 study of dostarlimab, an anti-PD-1 mAb, in 12 patients with mismatch repair-deficient stage II or III rectal adenocarcinoma, and found all patients studied had a clinical complete response, with no evidence of tumor on magnetic resonance imaging, positron-emission tomography, biopsy, etc., and no progression or recurrence during 6 to 25 months follow-up study^7^. However, limitations such as high cost, low stability and no oral bioavailability related to mAbs still exist^4,5^.

A promising alternative for mAb therapy is small-molecule inhibitors targeting the PD-1/PD-L1 pathway^4,5,^ with a shorter biological half-life, less side effect, easier administration routes and usually a lower price. Since the complete complex structures of human PD-1 and PD-L1 proteins had been elucidated in 2015^8^, a growing number of studies have been focused on searching for small-molecule PD-1/PD-L1 inhibitors. The PD-1/PD-L1 natural product inhibitors, including macrocyclic, phenolic and heterocyclic compounds, were reported Bristol-Myers Squibb (BMS) company developed a series of ^9.^ Bristol-Myers Squibb (BMS) company developed a series of macrocyclic peptides targeting PD-L1^4^, and also a series of small-molecule inhibitors of PD-1/PD-L1 with very low IC_50_ values in nM to μM range, whose structures had scaffolds based on substituted biphenyl group connected to another aromatic ring via a benzyl ether bond^4, 10^. The X-ray crystallographic study demonstrated that the BMS-202/PD-L1 complex structure is an asymmetric unit of four molecules of PD-L1 organized into two dimers, with each dimer bound to a single BMS-202 molecule^10^. Incyte corporation reported several PD-1/PD-L1 small molecule inhibitors with IC_50_ values in nM to μM range according to homogenous time-resolved fluorescence (HTRF) assay^4^, whose structures had fused heteroaromatic rings containing nitrogen atoms connected to a biphenyl moiety. Recently, novel benzo[c][1,2,5]oxadiazole derivatives have been designed and synthesized as potent small molecule inhibitors targeting the PD-1/PD-L1 pathway^11^. Among large number of small-molecule and peptide inhibitors of PD-1/PD-L1, small-molecule CA-170 and macrocyclic peptide BMS-986189 proceeded to clinical trial^4^. Searching for new compounds with PD-1/PD-L1 inhibitory activity requires a substantial amount of time, cost and effort. Drug repurposing, a strategy to identify new use of existing medications, can overcome these obstacles due to its low risk of failure, confirmed clinical safety and short study time^12^. One example is pyrvinium, an FDA approved anthelmintic drug, showed the PD-1/PD-L1 inhibition with an IC_50_ value of about29.66 μM^13^. Dietary polyphenol resveratrol (RSV) was reported to directly target PD-L1 glycosylation and dimerization and raise strong exacerbation of the cytotoxic activity of T-cells against breast cancer cells^14^. Molecular docking and molecular dynamics (MD) simulations predicted RSV to occupy the target space of BMS-202, which was shown to bind to PD-L1 and induced its dimerization^14^. On the other hand, it is worth noting that endogenous metabolites form a large group of small molecules, among which are some bioactive compounds that can regulate immune cell function, tumor progression and metastasis^15, 16^. For example, kynurenine induce regulatory T cell (T_reg_) cells through activation of the aryl hydrocarbon receptor^16^, and the antioxidant N-acetylcysteine and vitamin E were reported to stabilize BACH1 that makes cells more prone to migration, promoting the metastasis of cancer cells^17^. Searching for bioactive endogenous compounds is attractive, since most of them have significantly low side effects and with low cost. Up to now, there have been few studies on endogenous small molecule regulators of PD-L1 activity. Yuan et al. screened a home-built metabolic compound library and found pyridoxal, a member of vitamin B6, exerted prominent suppression effect on PD-L1 expression in the SW1990 and HepG2 cells upon treatment of 100, 200 and 400 μM for 24 h, by accelerating PD-L1 degradation in a proteasome-dependent manner^12^. To our knowledge, there has not yet any study on discovery of endogenous bioactive compounds with direct PD-1/PD-L1 inhibition. Considering various self-regulation processes in human body, we hypothesize that there may be some endogenous compound as PD-L1 inhibitor under physiological conditions.

Due to above mentioned characteristic structures of small-molecule PD-L1 inhibitors and the aim to target the binding between PD-1 and PD-L1, we selected potential cancer related metabolites with phenyl or heterocyclic ring moieties for endogenous small-molecule PD-L1 inhibitor screening. The first group of endogenous compounds includes tryptophan metabolites which have indole or phenyl groups and were identified as discriminant metabolites in multiple tumor types^18-20^. The second group of endogenous compounds consist of adenosine, adenosine 5′-triphosphate (ATP) and cyclic adenosine monophosphate (cAMP). Adenosine was found to accumulate in tumor microenvironment and serve as a mediator in CD8^+^ T cell suppression^21^. Extracellular ATP was reported to influence tumor growth and tumor-host interactions in different ways: low-level ATP may promote tumor cell proliferation and immunosuppression, while high-level ATP may promote antitumor immunity and activate infiltrating inflammatory cells^22^. ATP can also lead to immunosuppression by being degraded to adenosine^22^, and one critical mechanism of cancer immune evasion is the generation of high-level immunosuppressive adenosine in tumor microenvironment^23^. cAMP is synthesized from intracellular ATP by adenylyl cyclase (AC). In most tumor cells, increasing intracellular concentrations of cAMP can arrest cancer cell growth, induce apoptosis and limit cell migration^24^. cAMP was found to be significantly decreased in abundance in extracellular vesicles from prostate cancer patients with perineural invasion compared to those with no invasion^25^. The third group of endogenous compounds includes trace amines, which can bind to trace amine receptor 1 (TAAR1). Upon stimulation, TAAR1 triggers the accumulation of intracellular cAMP via AC activation and stimulates inwardly rectifying K^+^ channels^26^. Increased TAAR1 expression was found to be associated longer overall survival of breast cancer patients^27^.

The conventional approach for bioactive component screening requires bioassay-guided isolation, which is labor-intensive and time consuming with relatively high risk of failure due to irreversibly adsorption, decomposition and dilution effects, and sometimes may lead to false positives or negatives^28^. In recent years, bio-affinity ultrafiltration (BAU) coupled with ultraperformance liquid chromatography - high resolution mass spectrometry (UPLC-HRMS) has been developed to fulfill the purpose of rapid screening and identification of compounds that are potential ligands of target proteins. In BAU, the ligand-receptor complexes and unbound ligands are separated, and after being released from the complexes, the ligands are identified by UPLC-HRMS. BAU is a label-free method that works in a solution-based system, which ensures proper protein folding and stability. It consumes only a small amount of purified biomolecular receptor (usually microgram quantities)^29^. Theoretically, molecular docking and molecular dynamics (MD) simulations play an important role in protein inhibitor discovery and they can provide insightful information on the geometric features and energies of the ligand binding to the active site(s) of the receptor^30^.

In this study, we first used BAU and UPLC-HRMS to search for the candidates with potential PD-L1 affinity from 22 pre-selected endogenous metabolic compounds. Then, the inhibitory activities of the candidate compounds on the PD-1/PD-L1 binding were assessed by HTRF assay. The direct binding between the inhibitory compound and PD-L1 was investigated by surface plasmon resonance (SPR) technique. Molecular docking and MD simulations were performed to validate the binding properties of the candidate compounds with PD-L1, which was compared with the binding of the PD-1/PD-L1 lead inhibitor BMS-202. The results showed that cAMP may exert direct inhibition on PD-1 and PD-L1 binding, and the IC_50_ value determined by HTRF assay was 2.7 µM.

## Results and discussion

### Selected Endogenous Compounds - PD-L1 docking

Since most small molecule PD-1/PD-L1 inhibitors have characteristic structures containing one or more aromatic or heteroaromatic ring with nitrogen atoms^4^, we selected 22 endogenous compounds for BAU screening based on two criteria: 1) with aromatic, five or six-membered heterocyclic aromatic ring or fused heterocyclic ring structure; 2) reported to be differential metabolites associated with cancer or have effects on tumor cell proliferation or migration. The PD-1/PD-L1 inhibitor BMS-202 was used as a positive control. The 2D structures of selected 22 compounds and positive control BMS-202 were shown in Figure S1. All the selected compounds were found to have docking scores less than -5.0, among which BMS-202, β-NAD, ATP, adenosine, and cAMP have docking scores of -10.35, -9.29, -9.28, - 7.18, and -7.2, respectively (Table 1).

**Table 1.**
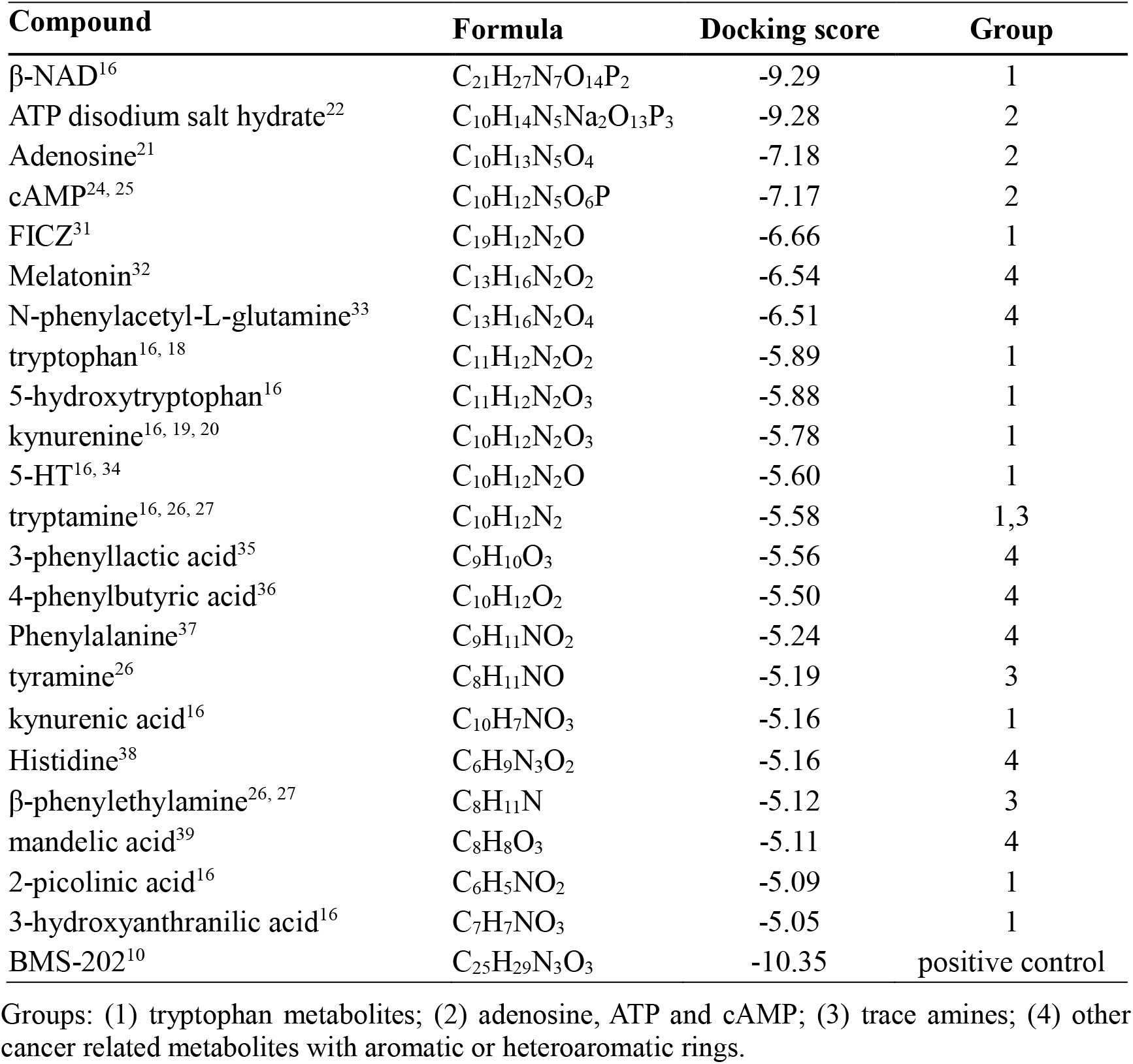
GBVI/WSA dG docking scores of selected 22 compounds binding to PD-L1.

| Compound | Formula | Docking score | Group |
| --- | --- | --- | --- |
| β-NAD<sup>16</sup> | C<sub>21</sub>H<sub>27</sub>N<sub>7</sub>O<sub>14</sub>P<sub>2</sub> | -9.29 | 1 |
| ATP disodium salt hydrate<sup>22</sup> | C<sub>10</sub>H<sub>14</sub>N<sub>5</sub>Na<sub>2</sub>O<sub>13</sub>P<sub>3</sub> | -9.28 | 2 |
| Adenosine<sup>21</sup> | C<sub>10</sub>H<sub>13</sub>N<sub>5</sub>O<sub>4</sub> | -7.18 | 2 |
| cAMP<sup>24, 25</sup> | C<sub>10</sub>H<sub>12</sub>N<sub>5</sub>O<sub>6</sub>P | -7.17 | 2 |
| FICZ<sup>31</sup> | C<sub>19</sub>H<sub>12</sub>N<sub>2</sub>O | -6.66 | 1 |
| Melatonin<sup>32</sup> | C<sub>13</sub>H<sub>16</sub>N<sub>2</sub>O<sub>2</sub> | -6.54 | 4 |
| N-phenylacetyl-L-glutamine<sup>33</sup> | C<sub>13</sub>H<sub>16</sub>N<sub>2</sub>O<sub>4</sub> | -6.51 | 4 |
| tryptophan<sup>16, 18</sup> | C<sub>11</sub>H<sub>12</sub>N<sub>2</sub>O<sub>2</sub> | -5.89 | 1 |
| 5-hydroxytryptophan<sup>16</sup> | C<sub>11</sub>H<sub>12</sub>N<sub>2</sub>O<sub>3</sub> | -5.88 | 1 |
| kynurenine<sup>16, 19, 20</sup> | C<sub>10</sub>H<sub>12</sub>N<sub>2</sub>O<sub>3</sub> | -5.78 | 1 |
| 5-HT<sup>16, 34</sup> | C<sub>10</sub>H<sub>12</sub>N<sub>2</sub>O | -5.60 | 1 |
| tryptamine<sup>16, 26, 27</sup> | C<sub>10</sub>H<sub>12</sub>N<sub>2</sub> | -5.58 | 1,3 |
| 3-phenyllactic acid<sup>35</sup> | C<sub>9</sub>H<sub>10</sub>O<sub>3</sub> | -5.56 | 4 |
| 4-phenylbutyric acid<sup>36</sup> | C<sub>10</sub>H<sub>12</sub>O<sub>2</sub> | -5.50 | 4 |
| Phenylalanine<sup>37</sup> | C<sub>9</sub>H<sub>11</sub>NO<sub>2</sub> | -5.24 | 4 |
| tyramine<sup>26</sup> | C<sub>8</sub>H<sub>11</sub>NO | -5.19 | 3 |
| kynurenic acid<sup>16</sup> | C<sub>10</sub>H<sub>7</sub>NO<sub>3</sub> | -5.16 | 1 |
| Histidine<sup>38</sup> | C<sub>6</sub>H<sub>9</sub>N<sub>3</sub>O<sub>2</sub> | -5.16 | 4 |
| β-phenylethylamine<sup>26, 27</sup> | C<sub>8</sub>H<sub>11</sub>N | -5.12 | 3 |
| mandelic acid<sup>39</sup> | C<sub>8</sub>H<sub>8</sub>O<sub>3</sub> | -5.11 | 4 |
| 2-picolinic acid<sup>16</sup> | C<sub>6</sub>H<sub>5</sub>NO<sub>2</sub> | -5.09 | 1 |
| 3-hydroxyanthranilic acid<sup>16</sup> | C<sub>7</sub>H<sub>7</sub>NO<sub>3</sub> | -5.05 | 1 |
| BMS-202<sup>10</sup> | C<sub>25</sub>H<sub>29</sub>N<sub>3</sub>O<sub>3</sub> | -10.35 | positive control |

Groups: (1) tryptophan metabolites; (2) adenosine, ATP and cAMP; (3) trace amines; (4) other cancer related metabolites with aromatic or heteroaromatic rings.

### Screening of potential endogenous small-molecule PD-L1 inhibitors with BAU-UPLC-HRMS method

In the BAU experiment, compounds were incubated with PD-L1 for 1.5 h to allow for binding of ligands to protein targets. Then the unbound compounds from the mixture were washed off, and the bound ligands were released and detected using UPLC-HRMS. By comparing the difference of compound EIC peak areas between experiments with normal and denatured PD-L1 proteins, candidate ligands in the mixture can be identified. The relative peak area enhancement of positive control BMS-202 was 23.59% (Figure S2), which was used as a standard to determine the candidate compounds bound to PD-L1. As a result, cAMP, tryptamine and 5-HT were found as candidate compounds binding to PD-L1, with relative peak area enhancements were 61.20, 40.98% and 60.94, respectively (Figure 1 and Table S1), and these three compounds were selected for subsequent activity test.

**Figure 1.**
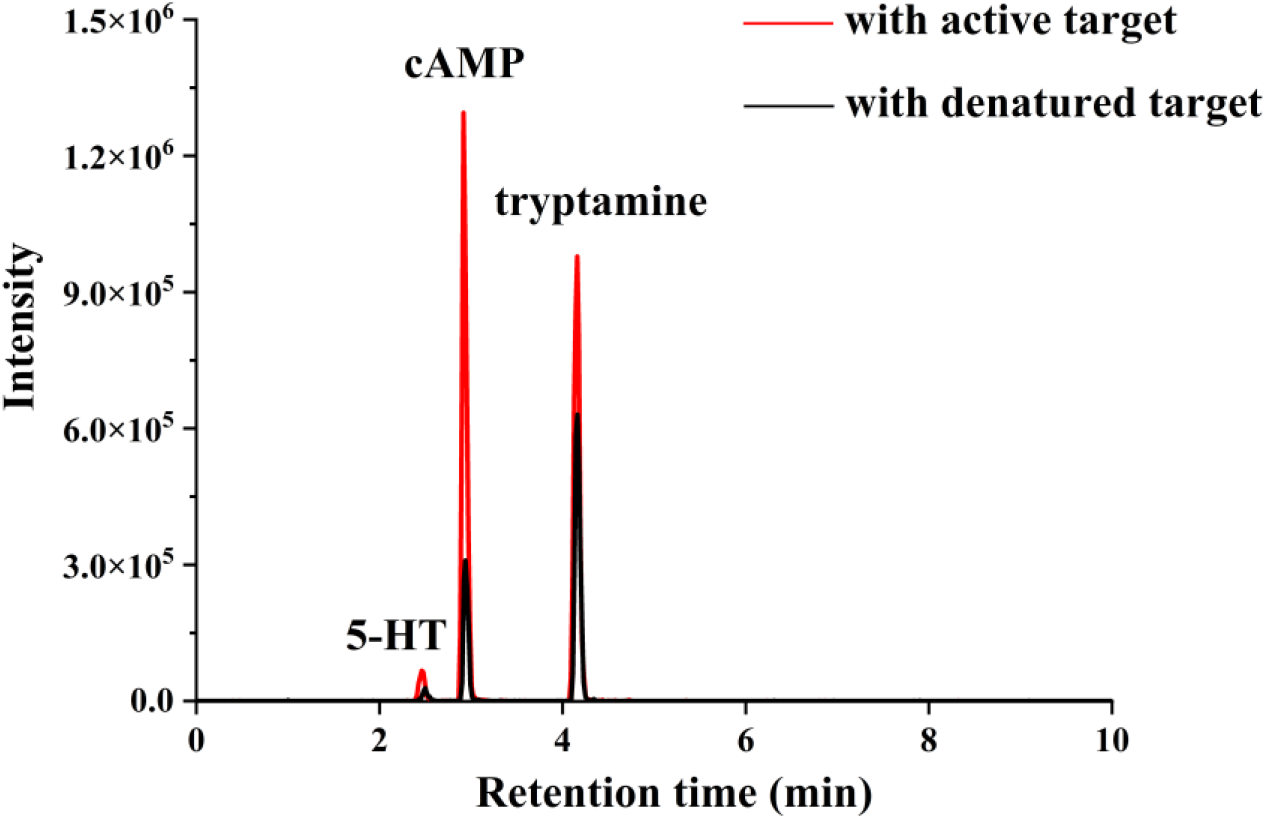
EIC of 5-HT (*m/z* = 177.1022 ± 0.005), cAMP (*m/z* = 330.0612 ± 0.005) and tryptamine (*m/z* = 161.1073 ± 0.005) in UPLC-HRMS analysis after the BAU experiment, n = 3.

### Inhibitory activity of candidate compounds against PD-1/PD-L1 interaction

In vitro HTRF assay was applied to assess the inhibitory activity of cAMP, tryptamine and 5-HT filtered out from the BAU experiment. Surprisingly, cAMP exhibited distinct inhibitory activity against the PD-1/PD-L1 interaction, with a half-maximal inhibitory concentration (IC_50_) of 2.7 µΜ (Figure 2A), whereas tryptamine and 5-HT did not show observable inhibition of PD-1/PD-L1 interaction in the concentration range of 1-100 µΜ (Figure S3). The IC_50_ of the positive control BMS-202 was derived as 11.4 nM (Figure 2B), which was the same order of magnitude with the reported value of 18 nM^40^.

**Figure 2.**
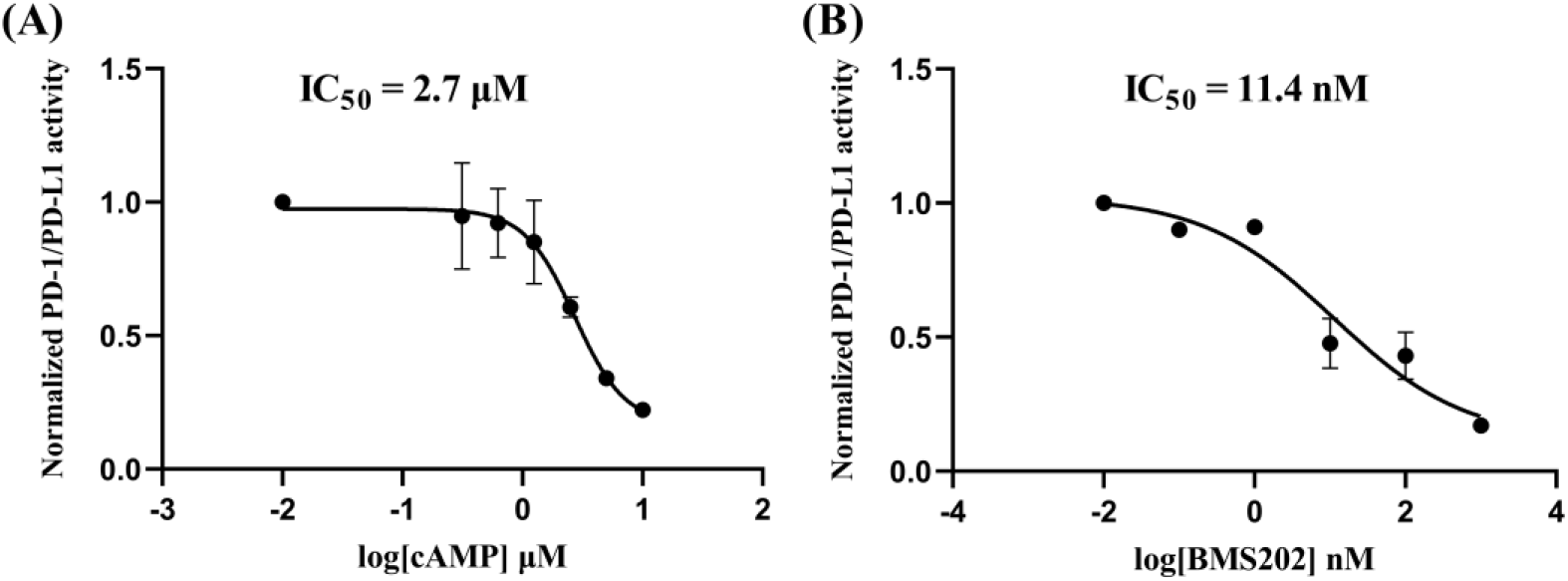
PD-1/PD-L1 inhibitory activity of (A) cAMP compared to (B) BMS-202 by HTRF assay. Data are mean ± standard deviation. cAMP n = 3, BMS-202 n = 2.

### Binding affinity of cAMP to PD-L1 by SPR analysis

SPR is a label free technique that can detect, monitor and quantitatively measure intermolecular interactions in real time^9^. We further performed SPR analysis to evaluate the binding affinity of cAMP to PD-L1. The equilibrium dissociation constant (K_D_) of cAMP was determined to be 1.72 mM by the saturation binding assay (Figure 3A), and K_D_ of BMS-202 was 36.73 µM (Figure 3B). The much larger K_D_ values of cAMP and BMS-202 compared with their IC_50_ values of 2.7 µΜ (cAMP) and 11.4 nM (BMS-202) determined from the HTRF assay may be ascribed to the fact that in the SPR experiment, PD-L1 proteins were fixed on the sensor chip surface restricting their fluidity and the K_D_ values obtained corresponded to the binding of cAMP or BMS-202 to monomeric PD-L1. However, the reported crystallographic structure demonstrated the binding mechanism between BMS-202 and PD-L1, in which BMS-202 induced the PD-L1 dimerization and bound to the PD-L1 dimer^10^, and we speculated cAMP might bind in the same way, which was subsequently validated using MD simulations. Nonetheless, the results of the SPR analyses suggested that cAMP and BMS-202 could also bind to monomeric PD-L1 in weak (cAMP) and moderate (BMS-202) interaction modes, respectively.

**Figure 3.**
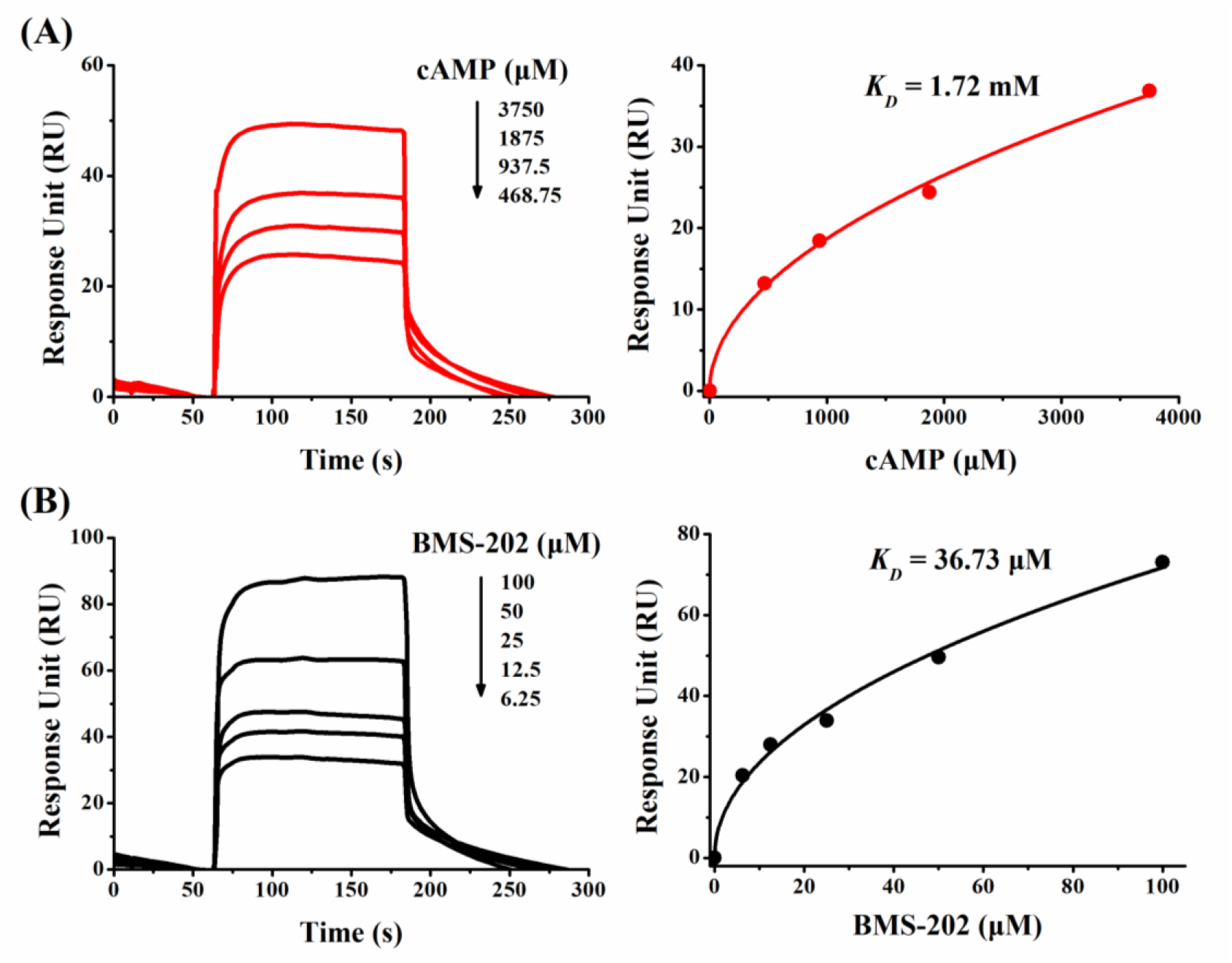
SPR sensorgrams by kinetic analysis of the binding of (A) cAMP and (B) BMS-202 to immobilized PD-L1 proteins. Plots of equilibrium responses versus compound concentrations were shown on the right, with the blank control response values subtracted from the experimental equilibrium response values. Estimated K_D_ values were 1.72 mM and 36.73 µM for cAMP and BMS-202, respectively.

### MD simulations on complexes of cAMP, 5-HT and tryptamine with dimeric PD-L1

To examine the binding properties of the candidate compounds with monomeric or dimeric PD-L1, 20 ns MD simulations were performed for the complexes of the three candidate endogenous compounds derived from the BAU experiment (cAMP, 5-HT and tryptamine), with PD-L1. The binding free energy of cAMP - dimeric PD-L1 was calculated to be -23.6 kcal/mol, lower than those of 5-HT (-20.9 kcal/mol) and tryptamine (-15.6 kcal/mol) (Table 2). The complex of BMS-202 with dimeric PD-L1 had lower binding free energy of -29.7 kcal/mol, consistent with the results of HTRF assay and the crystallographic structure of BMS-202 bound to PD-L1 reported in the literature^10^. In contrast, the binding free energy of the complex of cAMP - monomeric PD-L1 was -15.1 kcal/mol and that of cAMP - PD-1 was only -9.7 kcal/mol, indicating that cAMP tends to bind to two PD-L1 proteins and induce the dimerization of PD-L1, and such an interaction mode is similar to that of BMS-202. The generation of cAMP is mediated by the activation of adenylyl cyclase (AC), which catalyses intracellular ATP to cAMP conversion. The major receptor of cAMP in mammals is protein kinase A (PKA), which has two regulatory (R) and two catalytic (C) subunits^41^. Upon binding of cAMP molecules at the R subunits, the C subunits were activated and released, leading to PKA-dependent phosphorylation^41^. We also performed MD simulation of cAMP binding to the structure of two R subunits of PKA (PKA-Rs). The MD results predicted that the binding free energy of the cAMP - PKA-Rs complex was -34.8 kcal/mol (Table 2), significantly lower than that of the cAMP - dimeric PD-L1 complex (-23.6 kcal/mol), which was expected since PKA is the known major receptor of cAMP.

**Table 2.** MM/PBSA simulated van der Waals, electrostatic, polar solvation and binding free energies in kcal/mol of cAMP complexes.

| complex | van der Waals energy | electrostatic energy | polar solvation energy | non-polar solvation energy | binding free energy |
| --- | --- | --- | --- | --- | --- |
| cAMP - PKA-Rs | -40.0 | -58.3 | 66.7 | -3.2 | -34.8 |
| BMS-202 - dimeric PD-L1 | -50.0 | -7.3 | 31.6 | -4.0 | -29.7 |
| cAMP - dimeric PD-L1 | -45.5 | -14.1 | 41.0 | -5.0 | -23.6 |
| 5-HT - dimeric PD-L1 | -31.9 | -10.1 | 23.5 | -2.4 | -20.9 |
| Tryptamine - dimeric PD-L1 | -28.3 | -18.5 | 33.6 | -2.4 | -15.6 |
| cAMP - monomeric PD-L1 | -16.1 | -20.4 | -2.0 | 23.4 | -15.1 |
| cAMP - PD-1 | -21.2 | -23.1 | -2.4 | 37.0 | -9.7 |

Root mean square deviation (RMSD) measures structural conformation deviation in MD simulations. The RMSD results showed that the equilibrium of the cAMP - dimeric PD-L1 complex was obtained at ∼ 1.2 ns and stabilized at ∼ 5 Å displacement compared with its initial position (Figure 4A). Root mean square fluctuation (RMSF) measures the deviation of each atom relative to its initial position in MD simulations, showing the apparent flexible regions of the complex between 120 to 130 residue index (Figure 4B), which was the binding site in the middle of the PD-L1 dimer. In comparison, we performed MD simulations for BMS-202 binding to dimeric PD-L1 and cAMP binding to its known effector, PKA-Rs. The results of BMS-202 - dimeric PD-L1 complex was very similar to those of cAMP - dimeric PD-L1 complex, with equilibrium obtained at ∼ 6 ns and stabilized at ∼ 1.8 Å displacement compared to its initial position (Figure 4C), and apparent flexible region between 120 to 130 residue index (Figure 4D), indicating the binding site at the interface of the PD-L1 dimer. The cAMP - PKA-Rs complex reached equilibrium at ∼ 7.5 ns, and stabilized at ∼ 2.2 Å (Figure 4C), which was also similar to the RMSD result of the cAMP - dimeric PD-L1 complex, and RMSF plot showed flexible region between 180 and 220 residue index. The above results revealed the possible binding of cAMP to dimeric PD-L1 and the system binding mechanism closely mimicked that of BMD-202 and dimeric PD-L1.

**Figure 4.**
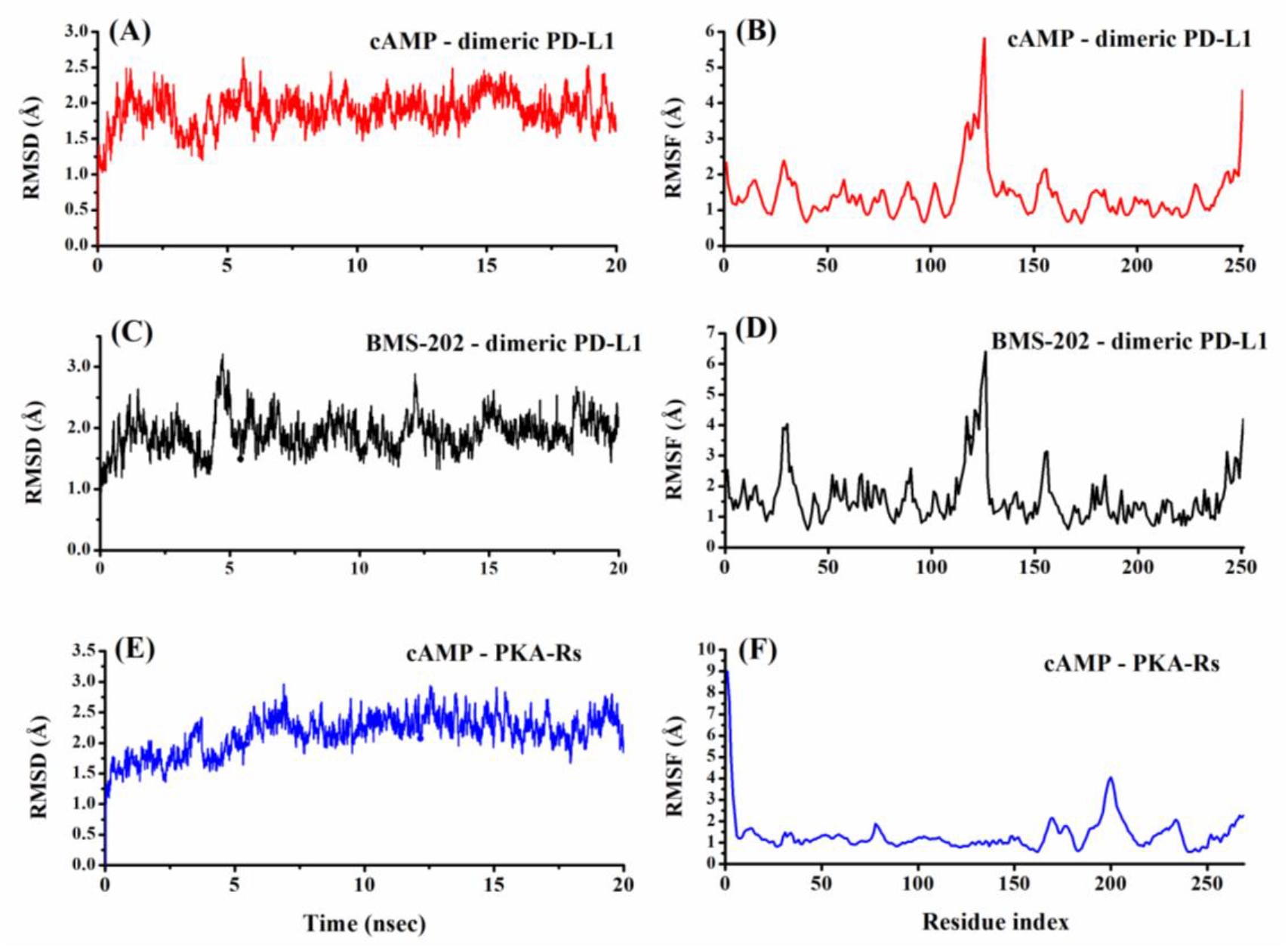
RMSD and RMSF of the cAMP - dimeric PD-L1 complex (A and B), BMS-202 - dimeric PD-L1 (C and D) and the cAMP - PKA-Rs complex (E and F).

### Binding properties between cAMP and dimeric PD-L1 predicted by molecular docking

According to the experimental results of HTRF, cAMP was shown to be capable of inhibiting PD-1/ PD-L1 interaction. To gain better understanding of the interacting modes between cAMP and PD-L1 or PD-1, molecular dockings were performed using the MOE package. Table 3 listed the docking scores of cAMP - dimeric PD-L1 (-7.17), cAMP - PKA-Rs (-7.80), cAMP - monomeric PD-L1 (-5.64), and BMS-202 with dimeric PD-L1 (-10.35). Therefore, cAMP is likely to bind to the dimerization surface of PD-L1, similar to BMS-202. Figure 5A showed the docking structure of _A_PD-L1/cAMP/ _B_PD-L1, in which cAMP located in a deep hydrophobic groove formed by the PD-L1 dimer. The cAMP - dimeric PD-L1 interactions included a hydrogen bond and an electrostatic interaction between _A_Lys124 and phosphoryl O atom of cAMP, a hydrogen bond between _A_Phe19 and hydroxyl O atom of cAMP, and one π-H and one π-π interaction between _A_Asp122/_B_Tyr56 and the 6-membered ring of cAMP, among which the cAMP electronic and hydrogen bonding interactions with _A_Lys124 was prominent. For BMS-202, there were one hydrogen bond between _A_Ala121 and imido group of BMS-202, one electrostatic interaction between _A_Asp122 and the N atom of five-membered nitrogen-containing heterocyclic ring of BMS-202, and two π-H interactions between _B_Met115/_B_Ala121 and two 6-membered rings of BMS-202, respectively, among which the hydrogen bond involved _A_Ala121 and the electrostatic interaction involved _A_Asp122 were dominant (Figure 5B). In the crystal structure reported by Zak et al.^10^, the central methyl-phenyl ring involved the hydrophobic interactions with _B_Met115 and the methoxy-pyridine moiety interacted with the side chain of _A_Asp122 via anion-π interaction, which were in general agreement with our findings. The dominant interactions between cAMP/BMS-202 and PD-L1 were both related to _A_Asp122. Compared to the BMS-202 - dimeric PD-L1, the cAMP - dimeric PD-L1 complex lacked interactions related to Ala121 and _B_Met115, but added interactions related to _A_Phe19 and _A_Lys124. In the PD-1/PD-L1 complex, the residues Lys124 and Asp122 were suggested to be involved in the strong electrostatic interactions, and Tyr56, Lys124 and Asp122 were considered to contribute to the hot regions of PD-L1, which may provide a basis for the discovery of new anti-PD-L1 small molecules^42^. Here, the interactions in relation to these three typical residues were present in both cAMP - dimeric PD-L1 and BMS-202 - dimeric PD-L1 complexes.

**Table 3.** GBVI/WSA dG docking scores of cAMP and BMS-202 with dimeric PD-L1, monomeric PD-L1, PD-1, and PKA regulatory subunits (PKA-Rs).

| **ligand** | **receptor** | **docking score** |
| --- | --- | --- |
| **cAMP** | dimeric PD-L1 | -7.17 |
|  | monomeric PD-L1 | -5.64 |
|  | PD-1 | -6.07 |
|  | PKA-Rs | -7.80 |
| **BMS-202** | dimeric PD-L1 | -10.35 |
|  | monomeric PD-L1 | -7.13 |
|  | PD-1 | -6.33 |
|  | PKA-Rs | -8.41 |

**Figure 5.**
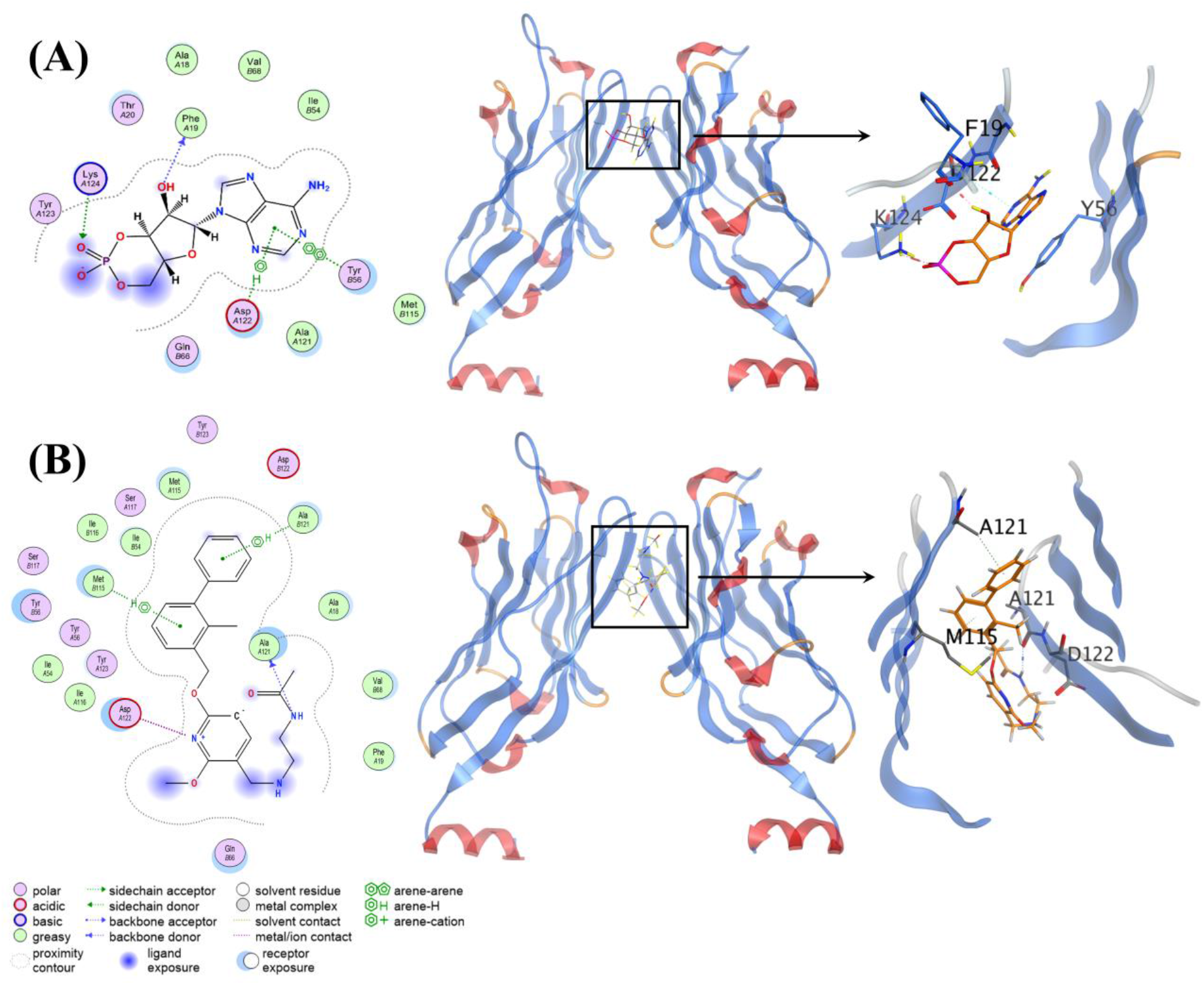
Detailed 2D and 3D representation of interactions between (A) cAMP and dimeric PD-L1 and (B) BMS-202 and dimeric PD-L1.

The docking results of cAMP and monomeric PD-L1 showed an interaction between Arg86 and hydroxyl O atom of cAMP, and two interactions between Arg86/Arg82 side chain and phosphoryl O atom/6-membered ring N atom of cAMP (Figure 6A). The docking results of cAMP with PD-1 showed an interaction between Ser93 and cAMP (Figure 6B). However, the Arg86, Arg82 and Ser93 residues were not hot regions involved in the PD-1 and PD-L1 binding^42^. Similarly, according to the docking scores and docking structures of BMS-202 - dimeric/monomeric PD-L1 and BMS-202 - PD-1 complexes (Table 3, Figure 5B, Figure S4A and S4B), BMS-202 was more likely to bind to dimeric PD-L1, in agreement with reported crystal structure^10^. Based on the docking results and MD simulations of the binding free energies, the inhibitory effect of cAMP on PD-1/PD-L1 activity was probably resulted from cAMP binding to dimeric PD-L1 other than monomeric PD-L1 or PD-1, with an interaction mode closely mimicking that of BMS-202. cAMP - PKA-Rs docking result showed interactions between Ala326/Arg333 and phosphoryl O atom of cAMP and Tyr371 accounted for hydrophobic interaction, which was expected since in the binding pocket of PKA regulatory domain, Ala326 was a hot spot amino acid in ligand binding, Tyr371 was responsible for hydrophobic interaction and Arg333 was involved in hydrogen bond interaction^43^.

**Figure 6.**
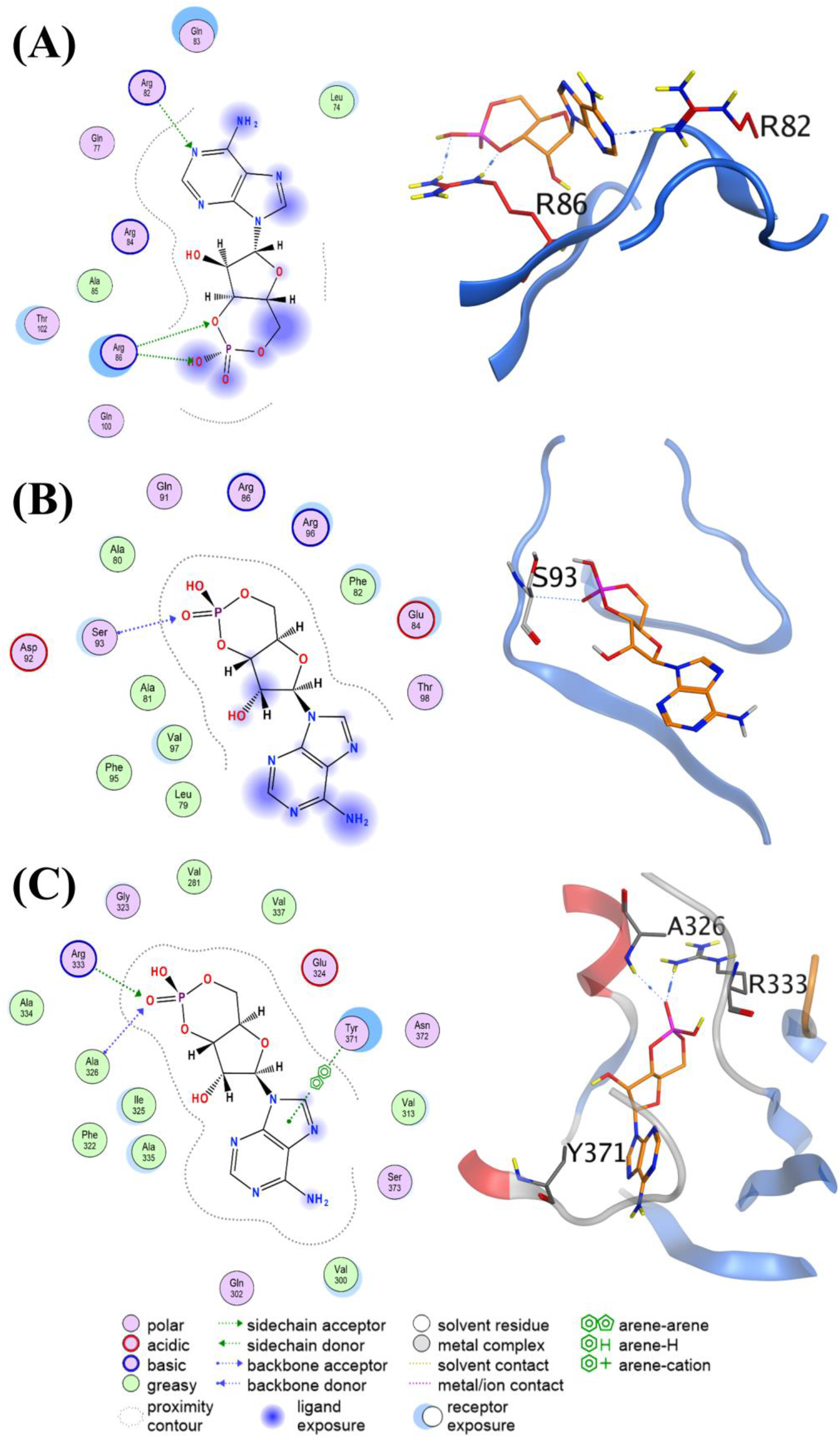
Detailed 2D and 3D representation of interactions between (A) cAMP and monomeric PD-L1, (B) cAMP and PD-1, (C) cAMP and PKA-Rs.

## Discussion

### cAMP as a PD-1/PD-L1 inhibitor by possibly binding to dimeric PD-L1

Small molecule inhibitors targeting PD-1/PD-L1 are attractive and urgently needed due to advantages such as fewer adverse side effects, less expensive than mAbs and possible oral bioavailablity^9^, but related research is still in early stage^44^. The endogenous compounds with aromatic and/or heterocyclic ring moieties and linked to tumorigenesis or anti-tumor activity might have some effects on the PD-1/PD-L1 pathway. In this work, we applied the BAU-UPLS-HRMS method combined with molecular docking and MD simulations to study the binding affinities of 22 selected endogenous compounds with PD-L1. Three compounds were identified to have potential PD-L1 affinity, among which cAMP showed a PD-1/PD-L1 inhibition with an IC_50_ value of 2.7 μM by HTRF analysis.

The binding interaction of cAMP - PD-L1 was further studied by the SPR assay. The K_D_ value of cAMP-PD-L1 binding was 1.72 mM, which was significantly larger compared with its IC_50_ value of 2.7 μM by the HTRF assay. Meanwhile, the SPR K_D_ value was shown to be 36 μM for BMS-202, which was also much larger than its IC_50_ value of 11.4 nM by HTRF assay. These large K_D_ values obtained from SPR analyses compared to the IC_50_ values determined from HTRF assays may be ascribed to the fact that in the SPR experiment, PD-L1 proteins were fixed on the surface of the sensor chip, thus the K_D_ values possibly corresponded to the ligand - monomeric PD-L1 binding. When PD-L1 was movable, BMS-202 induced and bound to dimeric PD-L1 as shown by the X-ray crystallography^10^. Similarly, cAMP would probably bind to dimeric PD-L1 other than monomeric PD-L1 in the PD-1/PD-L1 HTRF assay, resulting in a much smaller IC_50_ value of 2.7 μM.

Further, we performed MD simulations for the cAMP - dimeric or monomeric PD-L1 complexes. The binding free energies of the cAMP - dimeric and monomeric PD-L1 complexes were calculated to be about -23.6 and -15.1 kcal/mol, respectively, suggesting that cAMP was more likely to induce PD-L1 dimerization and bind to it. The RMSD data of the MD simulation of cAMP - dimeric PD-L1 system demonstrated its stability (Figure 4), and higher RMSF values at the binding site of cAMP - dimeric PD-L1 indicated that the binding site of the complex was flexible. At the binding interface of the cAMP - dimeric PD-L1 complex, _A_Lys124, _A_Phe19, _A_Asp122, and _B_Tyr56, which were involved in the electrostatic (_A_Lys124), hydrogen bonding (_A_Lys124 and _A_Phe19), π-H (_A_Asp122) and hydrophobic (_B_Tyr56) interactions, were residues related to the principal hot binding regions of PD-L1, and Tyr56 was reported to play a crucial role in the hydrophobic interactions in the BMS-202 - dimeric PD-L1 complex.

### cAMP regulation of PD-L1 expression and effects on immunocytes and cancer cells in the literature

PD-L1 expression is widely assumed to be induced by inflammatory cytokines, e.g, IFN-γ or TNF-α^45^, as well as oncogenic driver mutations^46^. The translation initiation factor eIF5B was demonstrated to be a key mediator of integrated stress-response-dependent translation of PD-L1 in lung cancer^45^, and the WW domain-containing transcriptional regulator 1 (TAZ) was shown to enhance PD-L1 levels in breast and lung cancer cell lines^47^.

Besides, several studies reported correlations between cAMP level and PD-L1 regulation. Sasi et al. reported that raising the level of intracellular cAMP induced PD-L1 expression in activated B-cell-like diffuse large B-cell lymphoma (ABC-DLBCL) cells, but not in germinal center B-cell-like (GCB) -DLBCL cells, and found that the cAMP-PKA-CREB pathway did not directly activate the PD-L1 promoter, instead, PD-L1 expression induction in ABC-DLBCL cells was mediated by the expression and secretion of cytokines driven by cAMP in a PKA-dependent manner^48^. Liu et al. reported antagonist treatment of adenosine A1 receptor (ADORA1) induced expression of PD-L1 via the cAMP/CREB/ATF3 pathway^49^. In contrast, activation of the lactate receptor GPR81 decreased intracellular cAMP level and inhibited PKA activity in lung cancer cells, resulting in activation of TAZ and its interaction with the transcription factor TEAD was essential for TAZ activation of PD-L1^50^. To sum up, the cAMP effects on PD-L1 expression were indirect and in a PKA-dependent manner, and the mechanisms are not well defined yet.

cAMP governs many important cellular processes and any malfunctions may lead to pathology^51^. cAMP has long been recognized as immunosuppressive, due to suppressing T cell activation and proliferation, and regulating certain cytokines, such as IFN-γ and TNF-α^52^. cAMP was reported to induce senescence in T cells^53^. However, cAMP has also been shown to promote T helper 17 (Th17) cell expansion, stimulating inflammation^54, 55^. Decreased level of cAMP in conventional DCs (cDCs) promoted Th2 cell differentiation^56^ but increased level of cAMP promoted induction of Th17^57^. cAMP-PKA-CREB signaling was involved in regulating a set of transcription factors that control cDC2 and cDC17 phenotypes and subsequent Th2 and Th17 bias^58^.

On the other hand, increased level of intracellular cAMP was reported to promote apoptosis and limit growth and migration of cancer cells^24^. The levels of cAMP within cells are controlled by ACs catalyzing cAMP synthesis and phosphodiesterases (PDEs) degrading cAMP. Marko et al. analyzed NCI panel of 60 human tumor cell lines and found high cAMP PDE activities in central nervous system, lung, breast and melanoma cell lines, and 41 of 60 cell lines with cAMP-specific PDE4 representing the highest AMP-hydrolyzing acitivity in the cytosol^59^. Treatment of PDE4 inhibitor DC-TA-46 induced cell cycle arrest in the G_1_/G_0_ phase and apoptotic cell morphology in all 7 human lung/breast tumor cell lines tested^59^. PDE4 inhibitors could cause a marked increase in intracellular cAMP levels, and mediated cancer cell growth suppression of HepG2 cells^24^, chemoresistant colon cancer^60^, pancreatic cancer cells^61^, brain tumor cells^62^, which has been suggested as a promising strategy for cancer treatment^24, 59^.

### cAMP targeting PD-1/PD-L1 interaction

From the discussion above, we can see that (1) cAMP effects on PD-L1 expression were considered to be indirect and not well defined yet, (2) cAMP can suppress T cell activation and proliferation, but can promote Th17 cell expansion, (3) cAMP can also suppress proliferation and migration of cancer cells. The immune regulation by cAMP is expected to be complicated and multifaceted. To our knowledge, there has been no study reported on whether cAMP can directly bind to PD-L1 proteins and exert any inhibitory effect on the PD-1/PD-L1 activity. By using BAU-LCMS method and PD-1/PD-L1 HTRF assay, we showed that cAMP is capable of inhibiting PD-1/PD-L1 activity, most likely by binding to and induce dimeric PD-L1. MD simulations predicted the capacity of cAMP to occupy the cleft formed by PD-L1 dimer in an interaction mode very similar to that of BMS-202. Thus, cAMP may play a role as a moderate PD-1/PD-L1 inhibitor by inducing PD-L1 dimerization. Our findings may broaden the understanding of PD-L1 regulation by cAMP.

### Comparison between the in vitro IC_50_ of cAMP on PD-1/PD-L1 binding and EC_50_ of cAMP on PKA

The basal cAMP concentration of cAMP was found to be about 50 - 150 nM in PC3, MCF7 and M628 cancer cells, less than 5 nM in CD4 cells, by measuring cytosolic cAMP in cell lysates using cAMP specific ELISA^53^, and ∼10 pmol/mg protein in human corpus cavernous smooth muscle^63^. The relatively high basal cAMP levels were reported to be around 1.2 μM and 1 μM in adult ventricular myocytes^64^ and CHO cells^65^, respectively, using FRET sensors. The different intracellular cAMP levels measured may be due to differences of cell types and methodological limitations. In addition, the in cell determination the apparent activation constant (EC_50_) of cAMP binding to PKA was measured to be 5.2 μM by a PKA-based FRET sensor, obviously higher than 0.3 μM measured in vitro^65^. In our study, the in vitro IC_50_ of cAMP suppressing the PD-1/PD-L1 interaction was shown to be about 2.7 μM, larger than the in vitro EC_50_ value (0.3 μM) of cAMP binding to PKA^65^. The in cell IC_50_ of cAMP targeting the PD-1/PD-L1 pathway may be expected to be larger than in vitro IC_50_, which is worth of further study.

Recently, Wang et al. reported a discovery of a PD-1/PD-L1 small molecular inhibitor, which induced the dimerization of PD-L1, which was internalized into the cytosol, and degraded through a lysosome-dependent pathway^66^. From this point of view, inducing the formation of PD-L1 dimer would be a good strategy to reduce the concentration of PD-L1 and inhibit PD-1/PD-L1 pathway. In this study, the endogenous compound cAMP showed a potential inhibitory effect on the PD-1/PD-L1 interaction, and it may induce and bind to dimeric PD-L1. Since the intracellular levels of cAMP can be controlled pharmacologically, it may provide an idea for the combined immunotherapy of cancer in the future.

## Supporting Information

**Materials and Methods** include detailed information on materials and reagents, BAU experiment, UPLC-HRMS analysis, HTRF binding assay, surface plasmon resonance (SPR) analysis, molecular docking and MD simulation methods.

**Table S1**. Relative peak area enhancement of selected compounds and positive control in BAU-UPLC-HRMS experiment.

**Figure S1**. Structure of selected 22 endogenous compounds and positive control BMS-202.

**Figure S2**. Relevant area enhancement of the positive control BMS202 (*m/z* = 420.2280 ± 0.005, ΔP = 23.59%).

**Figure S3**. Effects of 5-HT and trytamine on the PD-1/PD-L1 binding.

**Figure S4**. Detailed 2D and 3D representation of interactions between BMS-202 with (A) monomeric PD-L1, (B) PD-1, and (C) PKA-Rs.

A two-tailed unpaired Student’s t test was used for statistical analyses.

## Acknowledgements

This work was supported by Natural Science Foundation of Shandong Province (Grant No. ZR201911100165), and Laboratory of Marine Drugs and Biological Products, National Laboratory for Marine Science and Technology (Qingdao), 2019 Innovative talent introduction program (LMDBCXRC201904).

We would like to thank Xiaomin Zhang (Ocean University of China) for help with SPR experiments, and Xue Qiu (Ocean University of China) for offering counselling on HTRF assay.

## TOC

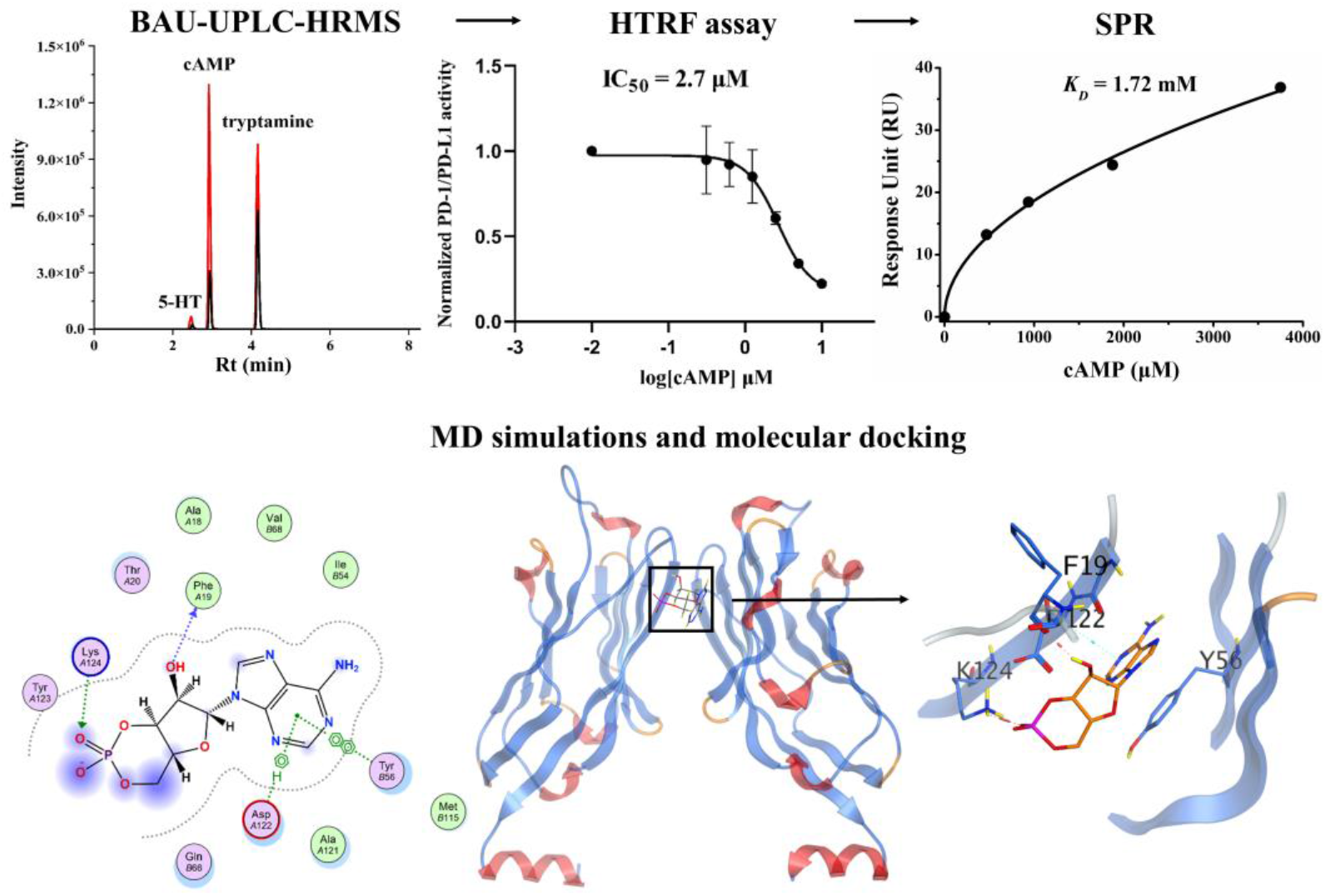

Synopsis: cAMP was found to directly inhibit PD-1/PD-L1 interaction using a combination of BAU-UPLC-HRMS, HTFR and SPR methods, via binding with dimeric PD-L1, verified by MD simulations and molecular docking.

## References

(1) He, X.; Xu, C. Immune checkpoint signaling and cancer immunotherapy. Cell research 2020, 30 (8), 660–669.

(2) Qin, W.; Hu, L.; Zhang, X.; Jiang, S.; Li, J.; Zhang, Z.; Wang, X. The Diverse Function of PD-1/PD-L Pathway Beyond Cancer. Frontiers in immunology 2019, 10, 2298.

(3) Wu, Y.; Chen, W.; Xu, Z. P.; Gu, W. PD-L1 Distribution and Perspective for Cancer Immunotherapy-Blockade, Knockdown, or Inhibition. Frontiers in immunology 2019, 10, 2022.

(4) Guzik, K.; Tomala, M.; Muszak, D.; Konieczny, M.; Hec, A.; Blaszkiewicz, U.; Pustula, M.; Butera, R.; Domling, A.; Holak, T. A. Development of the Inhibitors that Target the PD-1/PD-L1 Interaction-A Brief Look at Progress on Small Molecules, Peptides and Macrocycles. Molecules 2019, 24 (11), 2071.

(5) Wang, Y.; Guo, H.; Feng, Z.; Wang, S.; Wang, Y.; He, Q.; Li, G.; Lin, W.; Xie, X. Q.; Lin, Z. PD-1-Targeted Discovery of Peptide Inhibitors by Virtual Screening, Molecular Dynamics Simulation, and Surface Plasmon Resonance. Molecules 2019, 24 (20), 3784.

(6) Kim, E. S. Avelumab: First Global Approval. Drugs 2017, 77 (8), 929–937.

(7) Cercek, A.; Lumish, M.; Sinopoli, J.; Weiss, J.; Shia, J.; Lamendola-Essel, M.; El Dika, I. H.; Segal, N.; Shcherba, M.; Sugarman, R.; et al. PD-1 Blockade in Mismatch Repair-Deficient, Locally Advanced Rectal Cancer. The New England journal of medicine 2022.

(8) Zak, K. M.; Kitel, R.; Przetocka, S.; Golik, P.; Guzik, K.; Musielak, B.; Domling, A.; Dubin, G.; Holak, T. A. Structure of the Complex of Human Programmed Death 1, PD-1, and Its Ligand PD-L1. Structure 2015, 23 (12), 2341–2348.

(9) Liu, C.; Seeram, N. P.; Ma, H. Small molecule inhibitors against PD-1/PD-L1 immune checkpoints and current methodologies for their development: a review. Cancer cell international 2021, 21 (1), 239.

(10) Zak, K. M.; Grudnik, P.; Guzik, K.; Zieba, B. J.; Musielak, B.; Domling, A.; Dubin, G.; Holak, T. A. Structural basis for small molecule targeting of the programmed death ligand 1 (PD-L1). Oncotarget 2016, 7 (21), 30323–30335.

(11) Liu, L.; Yao, Z.; Wang, S.; Xie, T.; Wu, G.; Zhang, H.; Zhang, P.; Wu, Y.; Yuan, H.; Sun, H. Syntheses, Biological Evaluations, and Mechanistic Studies of Benzo[c][1,2,5]oxadiazole Derivatives as Potent PD-L1 Inhibitors with In Vivo Antitumor Activity. Journal of medicinal chemistry 2021, 64, 8391–8409.

(12) Yuan, J.; Li, J.; Shang, M.; Fu, Y.; Wang, T. Identification of vitamin B6 as a PD-L1 suppressor and an adjuvant for cancer immunotherapy. Biochemical and biophysical research communications 2021, 561, 187–194.

(13) Fattakhova, E.; Hofer, J.; DiFlumeri, J.; Cobb, M.; Dando, T.; Romisher, Z.; Wellington, J.; Oravic, M.; Radnoff, M.; Patil, S. P. Identification of the FDA-Approved Drug Pyrvinium as a Small-Molecule Inhibitor of the PD-1/PD-L1 Interaction. ChemMedChem 2021, 16 (18), 2769–2774.

(14) Verdura, S.; Cuyas, E.; Cortada, E.; Brunet, J.; Lopez-Bonet, E.; Martin-Castillo, B.; Bosch-Barrera, J.; Encinar, J. A.; Menendez, J. A. Resveratrol targets PD-L1 glycosylation and dimerization to enhance antitumor T-cell immunity. Aging 2020, 12 (1), 8–34.

(15) Geiger, R.; Rieckmann, J. C.; Wolf, T.; Basso, C.; Feng, Y.; Fuhrer, T.; Kogadeeva, M.; Picotti, P.; Meissner, F.; Mann, M.; et al. L-Arginine Modulates T Cell Metabolism and Enhances Survival and Anti-tumor Activity. Cell 2016, 167 (3), 829–842.

(16) Platten, M.; Nollen, E. A. A.; Rohrig, U. F.; Fallarino, F.; Opitz, C. A. Tryptophan metabolism as a common therapeutic target in cancer, neurodegeneration and beyond. Nature reviews. Drug discovery 2019, 18 (5), 379–401.

(17) Wiel, C.; Le Gal, K.; Ibrahim, M. X.; Jahangir, C. A.; Kashif, M.; Yao, H.; Ziegler, D. V.; Xu, X.; Ghosh, T.; Mondal, T.; et al. BACH1 Stabilization by Antioxidants Stimulates Lung Cancer Metastasis. Cell 2019, 178 (2), 330–345.

(18) Zang, X.; Monge, M. E.; Gaul, D. A.; Fernandez, F. M. Flow Injection-Traveling-Wave Ion Mobility-Mass Spectrometry for Prostate-Cancer Metabolomics. Analytical chemistry 2018, 90 (22), 13767–13774.

(19) Khan, A.; Choi, S. A.; Na, J.; Pamungkas, A. D.; Jung, K. J.; Jee, S. H.; Park, Y. H. Noninvasive Serum Metabolomic Profiling Reveals Elevated Kynurenine Pathway’s Metabolites in Humans with Prostate Cancer. Journal of proteome research 2019, 18 (4), 1532–1541.

(20) Mandarano, M.; Orecchini, E.; Bellezza, G.; Vannucci, J.; Ludovini, V.; Baglivo, S.; Tofanetti, F. R.; Chiari, R.; Loreti, E.; Puma, F.; et al. Kynurenine/Tryptophan Ratio as a Potential Blood-Based Biomarker in Non-Small Cell Lung Cancer. International journal of molecular sciences 2021, 22 (9), 4403.

(21) Mastelic-Gavillet, B.; Navarro Rodrigo, B.; Decombaz, L.; Wang, H.; Ercolano, G.; Ahmed, R.; Lozano, L. E.; Ianaro, A.; Derre, L.; Valerio, M.; et al. Adenosine mediates functional and metabolic suppression of peripheral and tumor-infiltrating CD8(+) T cells. Journal for immunotherapy of cancer 2019, 7 (1), 257.

(22) Vultaggio-Poma, V.; Sarti, A. C.; Di Virgilio, F. Extracellular ATP: A Feasible Target for Cancer Therapy. Cells 2020, 9 (11), 2496.

(23) Leone, R. D.; Emens, L. A. Targeting adenosine for cancer immunotherapy. Journal for immunotherapy of cancer 2018, 6 (1), 57.

(24) Massimi, M.; Ragusa, F.; Cardarelli, S.; Giorgi, M. Targeting Cyclic AMP Signalling in Hepatocellular Carcinoma. Cells 2019, 8 (12), 1511.

(25) Clos-Garcia, M.; Loizaga-Iriarte, A.; Zuniga-Garcia, P.; Sanchez-Mosquera, P.; Rosa Cortazar, A.; Gonzalez, E.; Torrano, V.; Alonso, C.; Perez-Cormenzana, M.; Ugalde-Olano, A.; et al. Metabolic alterations in urine extracellular vesicles are associated to prostate cancer pathogenesis and progression. Journal of extracellular vesicles 2018, 7 (1), 1470442.

(26) Pei, Y.; Asif-Malik, A.; Canales, J. J. Trace Amines and the Trace Amine-Associated Receptor 1: Pharmacology, Neurochemistry, and Clinical Implications. Frontiers in neuroscience 2016, 10, 148.

(27) Vattai, A.; Akyol, E.; Kuhn, C.; Hofmann, S.; Heidegger, H.; von Koch, F.; Hermelink, K.; Wuerstlein, R.; Harbeck, N.; Mayr, D.; et al. Increased trace amine-associated receptor 1 (TAAR1) expression is associated with a positive survival rate in patients with breast cancer. Journal of cancer research and clinical oncology 2017, 143 (9), 1637–1647.

(28) Chen, G.; Guo, M. Rapid Screening for alpha-Glucosidase Inhibitors from Gymnema sylvestre by Affinity Ultrafiltration-HPLC-MS. Frontiers in pharmacology 2017, 8, 228.

(29) Huang, Q.; Tang, J.; Chai, X.; Ren, W.; Wang, J.; Gan, Q.; Shi, J.; Wang, M.; Yang, S.; Liu, J.; et al. Affinity ultrafiltration and UPLC-HR-Orbitrap-MS based screening of thrombin-targeted small molecules with anticoagulation activity from Poecilobdella manillensis. Journal of chromatography. B, Analytical technologies in the biomedical and life sciences 2021, 1178, 122822.

(30) Naz, S.; Farooq, U.; Khan, S.; Sarwar, R.; Mabkhot, Y. N.; Saeed, M.; Alsayari, A.; Muhsinah, A. B.; Ul-Haq, Z. Pharmacophore model-based virtual screening, docking, biological evaluation and molecular dynamics simulations for inhibitors discovery against α-tryptophan synthase from Mycobacterium tuberculosis. Journal of Biomolecular Structure and Dynamics 2020, 39 (2), 610–620.

(31) Rannug, A.; Rannug, U. The tryptophan derivative 6-formylindolo[3,2-b]carbazole, FICZ, a dynamic mediator of endogenous aryl hydrocarbon receptor signaling, balances cell growth and differentiation. Critical reviews in toxicology 2018, 48 (7), 555–574.

(32) Talib, W. H. Melatonin and Cancer Hallmarks. Molecules 2018, 23 (3), 518.

(33) Zang, X.; Jones, C. M.; Long, T. Q.; Monge, M. E.; Zhou, M.; Walker, L. D.; Mezencev, R.; Gray, A.; McDonald, J. F.; Fernandez, F. M. Feasibility of detecting prostate cancer by ultraperformance liquid chromatography-mass spectrometry serum metabolomics. Journal of proteome research 2014, 13 (7), 3444–3454.

(34) Fatima, S.; Shi, X.; Lin, Z.; Chen, G. Q.; Pan, X. H.; Wu, J. C.; Ho, J. W.; Lee, N. P.; Gao, H. ; Zhang, G.; et al. 5-Hydroxytryptamine promotes hepatocellular carcinoma proliferation by influencing beta-catenin. Molecular oncology 2016, 10 (2), 195–212.

(35) Liu, J.; Geng, W.; Sun, H.; Liu, C.; Huang, F.; Cao, J.; Xia, L.; Zhao, H.; Zhai, J.; Li, Q.; et al. Integrative metabolomic characterisation identifies altered portal vein serum metabolome contributing to human hepatocellular carcinoma phyenyllactic acid biomarker. Gut 2021, 0, 1–11.

(36) Marrazzo, A.; Fiorito, J.; Zappala, L.; Prezzavento, O.; Ronsisvalle, S.; Pasquinucci, L.; Scoto, G. M.; Bernardini, R.; Ronsisvalle, G. Antiproliferative activity of phenylbutyrate ester of haloperidol metabolite II [(+/-)-MRJF4] in prostate cancer cells. European journal of medicinal chemistry 2011, 46 (1), 433–438.

(37) Ali, M. R.; Wu, Y.; Han, T.; Zang, X.; Xiao, H.; Tang, Y.; Wu, R.; Fernandez, F. M.; El-Sayed, M. A. Simultaneous Time-Dependent Surface-Enhanced Raman Spectroscopy, Metabolomics, and Proteomics Reveal Cancer Cell Death Mechanisms Associated with Gold Nanorod Photothermal Therapy. Journal of the American Chemical Society 2016, 138 (47), 15434–15442.

(38) Sirnio, P.; Vayrynen, J. P.; Klintrup, K.; Makela, J.; Karhu, T.; Herzig, K. H.; Minkkinen, I.; Makinen, M. J.; Karttunen, T. J.; Tuomisto, A. Alterations in serum amino-acid profile in the progression of colorectal cancer: associations with systemic inflammation, tumour stage and patient survival. British journal of cancer 2019, 120 (2), 238–246.

(39) Scheffer, D.; Kulcsar, G.; Czompoly, T. Identification of Further Components of an Anticancer Defense System Composed of Small Molecules Present in the Serum. Cancer biotherapy & radiopharmaceuticals 2019, 34 (3), 160–170.

(40) Chupak, L. S.; Zheng, X. Compounds useful as immunomodulators. WO2015034820 A1, 12 March 2015.

(41) Gold, M. G.; Lygren, B.; Dokurno, P.; Hoshi, N.; McConnachie, G.; Tasken, K.; Carlson, C. R.; Scott, J. D.; Barford, D. Molecular basis of AKAP specificity for PKA regulatory subunits. Molecular cell 2006, 24 (3), 383–395.

(42) Paciotti, R.; Agamennone, M.; Coletti, C.; Storchi, L. Characterization of PD-L1 binding sites by a combined FMO/GRID-DRY approach. Journal of computer-aided molecular design 2020, 34 (8), 897–914.

(43) Hameed, A.; Ashraf, S.; Israr Khan, M.; Hafizur, R. M.; Ul-Haq, Z. Protein kinase A-dependent insulinotropic effect of selected flavonoids. International journal of biological macromolecules 2018, 119, 149–156.

(44) Zhan, M. M.; Hu, X. Q.; Liu, X. X.; Ruan, B. F.; Xu, J.; Liao, C. From monoclonal antibodies to small molecules: the development of inhibitors targeting the PD-1/PD-L1 pathway. Drug discovery today 2016, 21 (6), 1027–1036.

(45) Suresh, S.; Chen, B.; Zhu, J.; Golden, R. J.; Lu, C.; Evers, B. M.; Novaresi, N.; Smith, B.; Zhan, X.; Schmid, V.; et al. eIF5B drives integrated stress response-dependent translation of PD-L1 in lung cancer. Nature cancer 2020, 1 (5), 533–545.

(46) Atefi, M.; Avramis, E.; Lassen, A.; Wong, D. J.; Robert, L.; Foulad, D.; Cerniglia, M.; Titz, B.; Chodon, T.; Graeber, T. G.; et al. Effects of MAPK and PI3K pathways on PD-L1 expression in melanoma. Clinical cancer research : an official journal of the American Association for Cancer Research 2014, 20 (13), 3446–3457.

(47) Janse van Rensburg, H. J.; Azad, T.; Ling, M.; Hao, Y.; Snetsinger, B.; Khanal, P.; Minassian, L. M.; Graham, C. H.; Rauh, M. J.; Yang, X. The Hippo Pathway Component TAZ Promotes Immune Evasion in Human Cancer through PD-L1. Cancer research 2018, 78 (6), 1457–1470.

(48) Sasi, B.; Ethiraj, P.; Myers, J.; Lin, A. P.; Jiang, S.; Qiu, Z.; Holder, K. N.; Aguiar, R. C. T. Regulation of PD-L1 expression is a novel facet of cyclic-AMP-mediated immunosuppression. Leukemia 2021, 35 (7), 1990–2001.

(49) Liu, H.; Kuang, X.; Zhang, Y.; Ye, Y.; Li, J.; Liang, L.; Xie, Z.; Weng, L.; Guo, J.; Li, H.; et al. ADORA1 Inhibition Promotes Tumor Immune Evasion by Regulating the ATF3-PD-L1 Axis. Cancer cell 2020, 37 (3), 324–339.

(50) Feng, J.; Yang, H.; Zhang, Y.; Wei, H.; Zhu, Z.; Zhu, B.; Yang, M.; Cao, W.; Wang, L.; Wu, Z. Tumor cell-derived lactate induces TAZ-dependent upregulation of PD-L1 through GPR81 in human lung cancer cells. Oncogene 2017, 36 (42), 5829–5839.

(51) Horvat, A.; Vardjan, N. Astroglial cAMP signalling in space and time. Neuroscience letters 2019, 689, 5–10.

(52) Mosenden, R.; Tasken, K. Cyclic AMP-mediated immune regulation--overview of mechanisms of action in T cells. Cellular signalling 2011, 23 (6), 1009–1016.

(53) Ye, J.; Ma, C.; Hsueh, E. C.; Dou, J.; Mo, W.; Liu, S.; Han, B.; Huang, Y.; Zhang, Y.; Varvares, M. A.; et al. TLR8 signaling enhances tumor immunity by preventing tumor-induced T-cell senescence. EMBO molecular medicine 2014, 6 (10), 1294–1311.

(54) Yao, C.; Sakata, D.; Esaki, Y.; Li, Y.; Matsuoka, T.; Kuroiwa, K.; Sugimoto, Y.; Narumiya, S. Prostaglandin E2-EP4 signaling promotes immune inflammation through Th1 cell differentiation and Th17 cell expansion. Nature medicine 2009, 15 (6), 633–640.

(55) Boniface, K.; Bak-Jensen, K. S.; Li, Y.; Blumenschein, W. M.; McGeachy, M. J.; McClanahan, T. K.; McKenzie, B. S.; Kastelein, R. A.; Cua, D. J.; de Waal Malefyt, R. Prostaglandin E2 regulates Th17 cell differentiation and function through cyclic AMP and EP2/EP4 receptor signaling. The Journal of experimental medicine 2009, 206 (3), 535–548.

(56) Lee, J.; Kim, T. H.; Murray, F.; Li, X.; Choi, S. S.; Broide, D. H.; Corr, M.; Lee, J.; Webster, N. J.; Insel, P. A.; et al. Cyclic AMP concentrations in dendritic cells induce and regulate Th2 immunity and allergic asthma. Proceedings of the National Academy of Sciences of the United States of America 2015, 112 (5), 1529–1534.

(57) Datta, S. K., Sabet, M., Nguyen, K. P., Valdez, P. A., Gonzalez-Navajas, J. M., Islam, S., Mihajlov, I., Fierer, J., Insel, P. A., Webster, N. J., Guiney, D. G., & Raz, E. Mucosal adjuvant activity of cholera toxin requires Th17 cells and protects against inhalation anthrax. PNAS 2010, 107, 10638–10643.

(58) Lee, J.; Zhang, J.; Chung, Y. J.; Kim, J. H.; Kook, C. M.; Gonzalez-Navajas, J. M.; Herdman, D. S.; Nurnberg, B.; Insel, P. A.; Corr, M.; et al. Inhibition of IRF4 in dendritic cells by PRR-independent and -dependent signals inhibit Th2 and promote Th17 responses. eLife 2020, 9:e49416.

(59) Marko, D.; Pahlke, G.; Merz, K. H.; Eisenbrand, G. Cyclic 3’,5’-nucleotide phosphodiesterases: potential targets for anticancer therapy. Chemical research in toxicology 2000, 13 (10), 944–948.

(60) McEwan, D. G.; Brunton, V. G.; Baillie, G. S.; Leslie, N. R.; Houslay, M. D.; Frame, M. C. Chemoresistant KM12C colon cancer cells are addicted to low cyclic AMP levels in a phosphodiesterase 4-regulated compartment via effects on phosphoinositide 3-kinase. Cancer research 2007, 67 (11), 5248–5257.

(61) Mouratidis, P. X.; Colston, K. W.; Bartlett, J. B.; Muller, G. W.; Man, H. W.; Stirling, D.; Dalgleish, A. G. Antiproliferative effects of CC-8062 and CC-8075 in pancreatic cancer cells. Pancreas 2009, 38 (1), 78–84.

(62) Goldhoff, P.; Warrington, N. M.; Limbrick, D. D., Jr.; Hope, A.; Woerner, B. M.; Jackson, E.; Perry, A.; Piwnica-Worms, D.; Rubin, J. B. Targeted inhibition of cyclic AMP phosphodiesterase-4 promotes brain tumor regression. Clinical cancer research : an official journal of the American Association for Cancer Research 2008, 14 (23), 7717–7725.

(63) Stief, C. G.; Uckert, S.; Becker, A. J.; Harringer, W.; Truss, M. C.; Forssmann, W. G.; Jonas, U. Effects of sildenafil on cAMP and cGMP levels in isolated human cavernous and cardiac tissue. Urology 2000, 55 (1), 146–150.

(64) Iancu, R. V.; Ramamurthy, G.; Warrier, S.; Nikolaev, V. O.; Lohse, M. J.; Jones, S. W.; Harvey, R. D. Cytoplasmic cAMP concentrations in intact cardiac myocytes. American journal of physiology. Cell physiology 2008, 295 (2), C414–422.

(65) Koschinski, A.; Zaccolo, M. Activation of PKA in cell requires higher concentration of cAMP than in vitro: implications for compartmentalization of cAMP signalling. Scientific reports 2017, 7 (1), 14090.

(66) Wang, T.; Cai, S.; Cheng, Y.; Zhang, W.; Wang, M.; Sun, H.; Guo, B.; Li, Z.; Xiao, Y.; Jiang, S. Discovery of Small-Molecule Inhibitors of the PD-1/PD-L1 Axis That Promote PD-L1 Internalization and Degradation. Journal of medicinal chemistry 2022, 65 (5), 3879–3893.

